# Expanding canonical cortical cell type markers in the era of single-cell transcriptomics

**DOI:** 10.1101/2025.08.26.672469

**Authors:** Dennis M Joshy, Soojin V Yi

**Affiliations:** Department of Mechanical Engineering, University of California, Santa Barbara, 93106, USA; Neuroscience Research Institute, University of California, Santa Barbara, 93106, USA; Ecology, Evolution, and Marine Biology, University of California, Santa Barbara, 93106, USA; Molecular, Cellular and Developmental Biology, University of California, Santa Barbara, 93106, USA

**Keywords:** scRNA-Seq, cell type markers, human cortex, sublayer specific markers

## Abstract

Cell type markers have been instrumental to physiological and molecular investigation of the human brain and remain essential for annotating cell type clusters in single-cell expression data and for targeted validation studies. However, expression of canonical markers in the target cell type (which we termed as the expression ‘fidelity’) as well as expression in unrelated cell types (which we termed as the ‘background expression’) across cortical regions remains poorly characterized. Using nearly 500,000 high-quality single-nucleus profiles from 19 studies, we quantified marker fidelity, revealing substantial regional variability. We developed a statistical framework that aggregates annotated barcodes into pseudo-bulk profiles, applied rigorous performance metrics, and identified markers with improved fidelity, reduced background, and consistent expression across regions. This approach extended the canonical marker set for six major brain cell types and yielded superior subtype-specific markers. The resulting marker lists, and a user-friendly analysis interface, provide a valuable resource for cell type annotation and validation in neurological research.

## Introduction

Identification of cell type in the central nervous system is crucial to understanding functional circuits in the brain and to developing translational products. Heterogeneous cell types in the brain have been traditionally recognized by their distinctive morphology, as well as gene expression profiles, leading to the catalogue of ‘cell-type markers’. These are especially well defined for major cell types, such as neurons, oligodendrocytes and oligodendrocyte precursor cells, astrocytes, and microglia (Mirsky; Kennedy).

Marker genes for these cell types tend to be proteins that are critical for the operation of the cell. For instance, some of the most well-known markers used for astrocytes are the Aquaporin-4 (*AQP4)* and Glial Fibrillary Acidic Protein (*GFAP*), which are essential for astrocyte function as the former is involved in facilitating water transport and ion homeostasis (Satoh et al.) and the latter being a major filament protein providing structural support (Lee et al.). *GAD1* and *GAD2,* which are central to the synthesis of GABA, are prominent inhibitory neuron markers (Westmoreland et al.), while the vesicular glutamate transporter SLC17A7 is commonly used to detect glutamatergic or generally excitatory neurons (Hodge et al.). The *MBP* (Zecevic et al.), *MOG* (Scolding et al.) and *OPALIN* (Golan et al.) genes are widely used to annotate oligodendrocyte cells. Other markers include Platelet-Derived Growth Factor Receptor Alpha (*PDGFRA*, (Takei et al.)) and protocadherin-15 (*PCDH15,* (Zhen et al.)) for oligodendrocyte precursor cells (OPCs, (Brenner et al.; Khrameeva et al.)). In microglial cells, Amyloid beta A4 precursor protein-binding family B member 1-interacting protein (*APBB1IP*) is an important marker involved in signal transduction, T-cell activation, and regulating cell adhesion (Pinosanu et al.). These genes will be hereafter referred to as ‘canonical cell type markers’.

Canonical cell type markers are foundational in the use of cell sorting to separate and isolate unique cell types (Gedye et al.; L. Ramos et al.; Berto et al.). They also provide complementary information in the era of single-cell studies. Single-cell RNA-Seq provides a comprehensive picture of cellular heterogeneity and allows us to capture combinatorial transcriptional signatures that would otherwise be averaged out in bulk RNA-Seq. They also facilitate a more nuanced and comprehensive understanding of cell types and states in the human brain, which is a key area of current research in neuroscience (e.g., (Darmanis et al.; Suresh et al.; Jorstad et al.; Adameyko et al.)). Nevertheless, the canonical cell type markers have been instrumental in annotating clusters of cells in many single-cell RNA-Seq experimental studies. In addition, canonical cell type markers are used for validation experiments, such as the use of *GFAP, AQP4* in the study of stroke (Weber et al.) and for evolutionary studies to identify astrocytes (Hodge et al.) in human and mouse motor cortex. *PDGFRA* and *MOG* have been used to distinguish between OPCs and oligodendrocytes in a comparison of human and nonhuman primate brains (Caglayan et al.).

As single-cell technology continues to become popular, techniques for identifying these markers are becoming increasingly integrated into analysis packages like Seurat (Butler et al.) and Scanpy (Wolf et al.). There are also multiple databases such as panglaoDB (Franzén et al.) and CellMarker (Hu et al.), and scBrainMap (Chi et al.). While these databases are great resources, only a few studies have examined the specificity of expression of these proposed markers in different brain regions. There are also recent work in biomarker discovery using single-cell data such as (McKenzie et al.) which introduced ideas of prioritizing specificity and enrichment and validated results. Other methods have relied on co-expression analyses (Qiu et al.). Even fewer studies have revisited many of the existing marker genes using transcriptomics and proteomics (Dai et al.)

The effectiveness of cell type markers to sort and isolate cells requires them to meet the following conditions: first, they should be expressed in the corresponding cell types, which we will refer to as *expression fidelity*, or simply, *fidelity*. Second, they should not be expressed or lowly expressed in non-corresponding cell types, which we will refer to as low *background expression*. These features are necessary to avoid potential mislabeling. Single-cell RNA-seq data provides an opportunity to examine these features at cellular resolution. Here, we were motivated by our observation that not all cells in a cell type cluster expressed the relevant canonical cell type markers in single-cell RNA-seq studies. In addition, some cells that are not within the corresponding clusters expressed marker genes; in other words, there was substantial background expression. These could be due to various reasons, such as variable cell state (due to cell cycle or other factors) or in some cases, technical artefacts in experimentation or sample collection or simply transcriptional noise and/or variability.

We thus aimed to utilize the available single-cell data to extend the widely used canonical marker set to discover additional novel marker genes that may perform equally well, if not better than the currently used canonical cell type markers in terms of expression fidelity and low background expression. In addition, it is standard practice to use multiple markers to fluorescently ascertain the identity of the cell. Therefore, there is a clear need to extend the current repertoire of cell type markers. In this study, we used 19 recent single cell RNA-seq datasets from 12 studies to propose a generalizable set of markers with comparable properties to canonical markers using pseudo-bulk aggregated statistical testing. We tune multiple parameters associated with this statistical approach and use performance evaluation metrics such as the Adjuster Rand Index and the Silhouette score. We further make this resource available to the community via an RShiny application. Our findings indicate that fidelity-based methods enhance our understanding of major and potentially layer-specific cell types in the human cortex.

## Results

### Traditional brain cell types exhibit varying canonical marker expression across regions

An integrative analysis of single-nucleotide RNA-seq datasets from different parts of the human cortex offers an opportunity to examine the expression levels of canonical cell type markers in distinct regions. We collected raw and processed single-cell count matrices from 19 recent datasets (Supplementary Table S1). The counts of barcodes following ambient RNA removal and single-cell QC (methods) are summarized in Supplementary Table S2. As described earlier, these studies used different sets of canonical cell type markers to annotate cells and to validate the performance of their annotation scheme. We therefore used these markers as a starting point or baseline for our methodology. From these studies, we used a set of 30 canonical cell type markers (Supplementary Table S3) that are used to classify six major cell types, namely excitatory and inhibitory neurons, oligodendrocytes, OPCs, microglia and astrocytes.

Post pseudo-bulk aggregation, we obtained a global matrix of 319 samples with 14,118 protein-coding genes common to all 19 datasets (simple intersection of aggregate matrices from each region). Principal component analysis (Methods, Supplementary Figure S1) shows that the normalized pseudo-bulk samples form clear clustering among cell types despite being obtained from various areas of the human cortex. We then examined the expression of the canonical marker set in these data sets. As expected, these markers yield clear clusters of major cell types (Figure 1A). However, the expression levels of canonical markers vary substantially across different regions (cluster-wise mean expression across corresponding cell type markers: 11.86 ± 1.55 S.D., Supplementary Table S3, at the pseudo-bulk level) Moreover, some markers exhibit variable levels of expression beyond the regional differences. Microglial cell markers like *AIF1* (S.D. 3.45) and oligodendrocyte markers like *PLP1* (S.D. 3.53), and *ST18* (S.D. 3.55) exhibit the highest variability.

**Figure 1.**
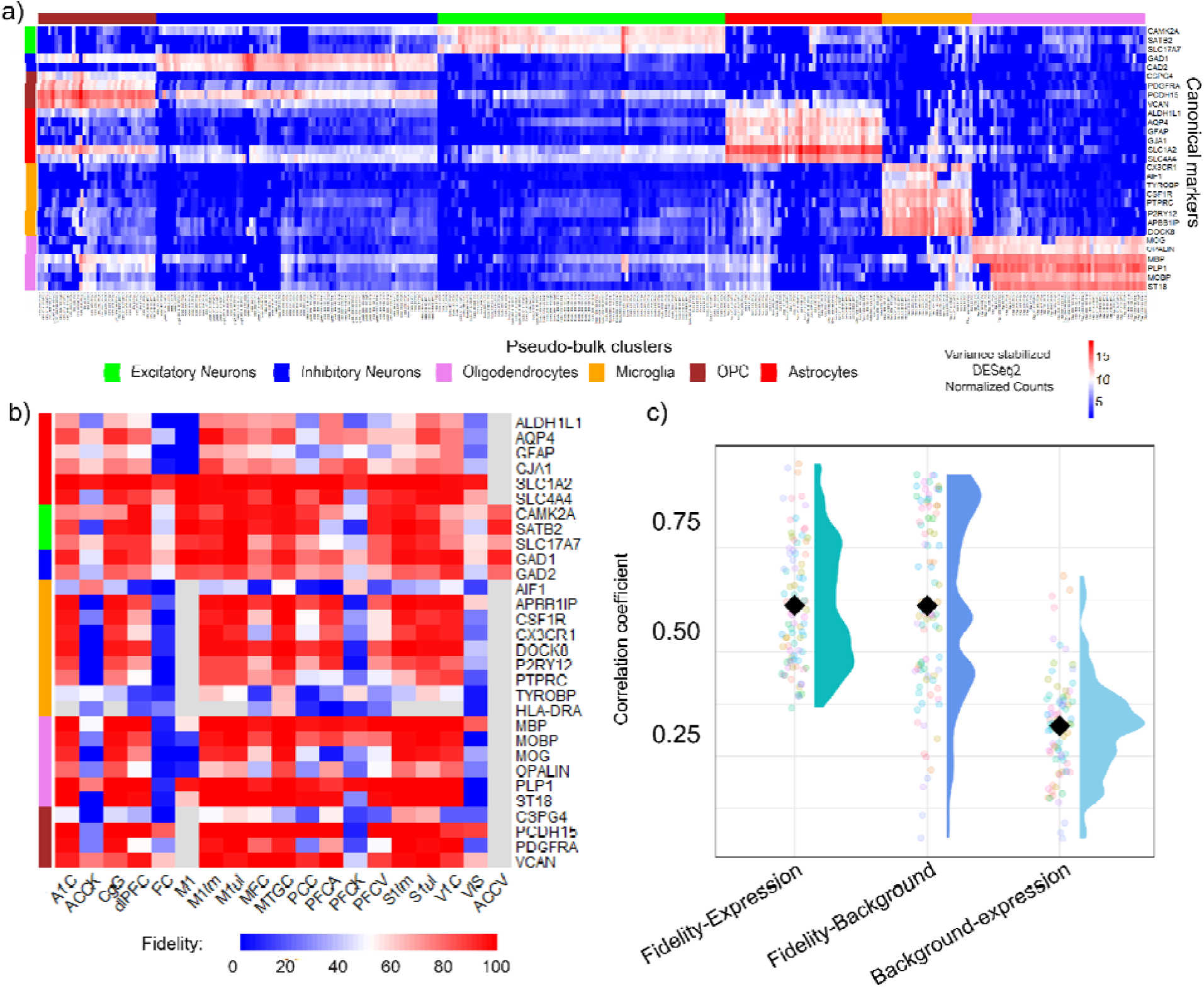
Canonical markers expression depends on cortical region, and the expressing proportion of cells is correlated with expression level. A) variance stabilized, normalized global expression matrix in heatmap form for plotted using all 319 pseudo-bulk clusters demonstrate varying levels of marker expression and potential background gene expression in some canonical markers in other cell types. variance stabilized, normalized global expression matrix in heatmap form for plotted using all 319 pseudo-bulk clusters, with canonical markers on the y-axis. B) Cell type cluster wise fidelity metrics calculated for each marker in each study (grayed cells indicate 0 cells in that cluster from that study). C) Correlation coefficients between fidelity, background and cluster-wise pseudo-bulk aggregated normalized counts (Black diamond represents mean of the distribution).

In addition, some markers show substantial levels of expression in other cell types (high background expression). As seen in Figure 1A, excitatory neuronal markers such as *CAMK2A* demonstrate background expression in inhibitory neurons. Well-known inhibitory neuron markers such as *GAD1* also appear to be upregulated in OPCs (brown). Widely used OPC markers such as *PCDH15* appear to be expressed in inhibitory neurons. High-expressing astrocyte markers such as *SLC1A2,* and *SLC4A4* also demonstrate expression in multiple cell-types including excitatory neurons, inhibitory neurons and OPCs. Oligodendrocyte markers such as *PLP1,* and *MBP* also appear to have noticeable expression in microglia and OPC cell types.

These results indicate that in addition to variability across different parts of the human cortex, canonical cell type markers exhibit variability across cell types, including low or no expression in corresponding cell types and substantial expression in non-corresponding cell types. To more formally quantify these aspects, we examined the fractions of cells in each corresponding cluster that express at least 1 transcript of the corresponding cell type marker. We define *expression fidelity* as the proportion of the cells in a cluster expressing the relevant canonical marker gene for that cell type (Figure 1B). The term *background* is defined as the number of cells outside this cluster of cells expressing canonical markers of the concerned cluster (Supplementary Figure S1). The heatmap in Figure 1B demonstrates substantial variability in the proportion of cells expressing these canonical cell type markers. Inhibitory neuron markers express the highest mean cluster-wise region-averaged fidelity of 77.4% + 1.41 S.D., while microglial cell type markers have the lowest value of 54.2% + 1.11 S.D (Figure 1B). Astrocyte cells demonstrate the highest variability in regional canonical marker fidelity with a standard deviation of 9.30 %. These results suggest that a large proportion of cells within the defined clusters do not necessarily express these canonical cell type markers.

We further examined the relationship between fidelity, background and the expression levels of genes in each cluster. Since fidelity and background are cluster-wise metrics, we pseudo-bulk aggregated by summing the counts from all samples for a given cell type. We then normalized this using DESeq2 and calculated the correlation coefficient between the fidelity and background signals for each gene for all samples (Figure 1C). From Figure 1C, we observe that fidelity and expression are correlated (average correlation coefficient: 0.61). The correlation is highest for microglia (average correlation coefficient: 0.747) and lowest for excitatory neurons (average correlation coefficient: 0.529). Similarly, we observe that the background and fidelity signals are also highly correlated (average correlation coefficient: 0.61), while the correlation between background and expression is lower (average correlation coefficient: 0.32). These results indicate that choosing high expressing genes can result in high background signals and that selecting marker genes requires careful consideration of both expression levels and marker fidelity and background.

### Fidelity informed marker selection yields novel cell type markers

Differential expression analysis provides us with the statistical support to identify biomarkers in single-cell data clusters. However, as seen above, this does not automatically translate to high expression fidelity within the corresponding cell type cluster. We hypothesized that explicitly incorporating thresholds of fidelity and background expression will yield novel, biologically relevant markers to efficiently distinguish cell types. We thus aimed to utilize this strategy to identify cell type markers that can reliably distinguish the six major cell types. We first performed a one-vs-all differential expression analysis for each cell vs all remaining 5 cell types using pseudo-bulk aggregated clusters. We merged all the DE results from the 19 datasets into one single table; we then selected genes that were significantly differential expressed with respect to the background (*Adjusted p-value < 0.05* and *log2FoldChange > 2)* and met the cell type specific thresholds of *fidelity* and *background*. These metrics ensured high cell type specificity locally in each study.

We examined the number of times that each gene in our study met these criteria out of all 19 studies and selected the top 5 genes that occurred the most frequently for each cell type. Furthermore, we implemented the *Cohen-d distance metric* to select markers that exhibit distinctive expression in the corresponding cell types in the global pseudo bulk matrix. The Cohen-d distance is used to calculate an effect size between the mean expression levels of a marker gene of one cell type vs all other cell types globally. We calculate the pooled standard deviation using the distribution of expression of a marker gene in the corresponding cell type pseudo-bulk clusters in all samples and studies and the corresponding distribution of expression of the remaining pseudo-bulk clusters in all studies. The Cohen’s d effect size is also used by the scran package for marker detection between clusters of cells and for meta-analyses across multiple studies (Lun et al.; Rajderkar et al.; Pullin and McCarthy). Additional details are provided in Materials and Methods.

Using this approach, we discovered a set of 24 markers that are reliably expressed in cell type specific manner in multiple brain regions and demonstrate high fidelity with low background expression (Figure 2A, B, Supplementary Figure S2). We will refer to this set as ‘calculated markers’ hereafter. Our approach also rediscovered some of the more robust canonical cell type markers including *GAD1, GAD2 (*Inhibitory Neuron*), SAT2B* (Excitatory Neuron)*, ST18, PLP1* (Oligodendrocytes), *CSF1R* (Microglia), *VCAN* (OPC), and *PDGFRA* (OPC). This high rate of overlap (*P*<10^-15^, hypergeometric test) emphasizes the utility of canonical cell type markers while supporting that our approach can identify biologically relevant markers. Supplementary Figure S3 demonstrates the background levels of these calculated markers.

**Figure 2:**
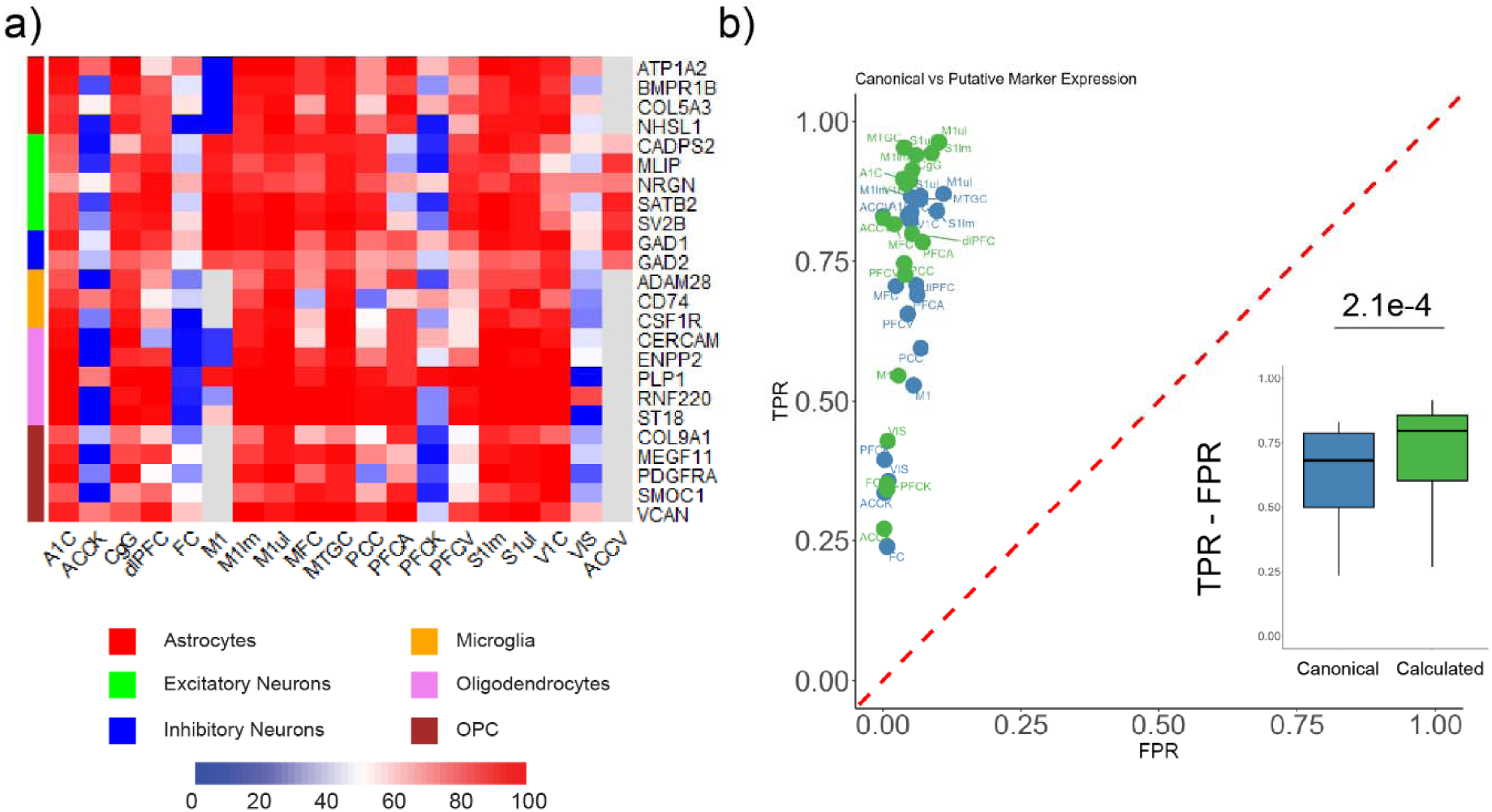
Calculated markers demonstrating increased fidelity, reduced background and higher AUROC scores. A) Calculated markers exhibit increased expression fidelities compared to the currently used canonical markers. B) ROC plotted for canonical and Calculated markers based on False Positive Rate (x-axis) and True Positive Rate (TPR - y-axis), with the line y=x for comparison. (Inset) calculated markers exhibit higher (TPR - FPR) than canonical markers (p-value = 2.1e-4).

The novel calculated markers include *NHSL1* and *ATP1A2* astrocyte genes. *NHSL1* is known to be involved in the regulation of cell migration and lamellipodia formation in multiple cell types (Law et al.). We further discovered microglial markers: *CD74* a marker for activated microglia and *ADAM28* which is enriched in activated microglia to regulate cytokine levels (Nuttall et al.). We also report calculated oligodendrocyte markers: *CERCAM,* a plasma membrane molecule involved in cell adhesion (Starzyk et al.). *RNF220* is another marker we identified that facilitates oligodendroglial development and myelination (Li et al.). We did not find additional inhibitory neuronal cell type markers of comparable expressive strength, fidelity and background as the reference *GAD1* and *GAD2* markers. However, we can tune our search parameters to identify genes like *GRIP2,* which is highly specifically expressed in inhibitory neurons. This may serve as a potential complementary inhibitory neuronal marker, with the caveat in mind that the fidelity levels of *GRIP2* are rather variable across different regions (Supplementary Figure S4). While our current report includes those selected under specific criteria, we realize that different groups would prefer to use markers based on different criteria according to their specific research design and goals. We have provided the above analysis and thresholding capability as an RShiny application as a resource for the community (https://dmj6288.shinyapps.io/CortexMapperV1/) to discover and test marker sets of various performance levels.

To statistically evaluate the performance of these markers, we applied the concepts of True Positive Rates (TPRs) and False Positive Rates (FPR) in terms of the number of cells that correctly express the corresponding markers. Expression fidelity and background correspond to TPR and FPR, as the former is the proportion of cells correctly expressing marker genes, and the latter correspond to the proportion of cells incorrectly expressing the marker genes. To visualize the effectiveness of the calculated and canonical marker sets, we visualize the trends on a Receiver-Over-Operator (ROC) curve (Figure 2B). We calculated the Area Under the ROC (AUROC) score and demonstrated that the calculated marker set has a 6.67% higher score (0.96) vs the canonical marker set (0.90). Comparing the distributions of the differences between the average TPR and the average FPR, we observed that calculated markers have a significantly higher difference (*P* < 2.1e10^-4^) than canonical cell type markers (Figure 2B, inset).

Using this set of 24 calculated markers we were able to obtain biologically meaningful clusters for all 6 cell types with a few outliers. We compared the clustering of canonical cell type markers and these calculated markers (comparing clustering in Figure 1A and Supplementary Figure S5) using the Adjusted Rand Index Score (ARI) and the Silhouette score, two widely used metrics to assess the quality of clustering (Hubert and Arabie; Rousseeuw; Kanter et al.; Peng et al.). The calculated markers-based clustering had a slightly improved pseudo-bulk ARI and Silhouette score (ARI : 0.995, Average Silhouette Score: 0.482) compared to that using canonical markers (ARI : 0.95, Average Silhouette Score: 0.44). These results indicate that the calculated markers provide a biologically relevant and improved distinction between major cell types.

### Calculated Marker set is generalizable across regions in classification and clustering tasks generalizable

To evaluate the ability of the calculated marker set to generalize across datasets in our meta-study, we performed Leave-One-Out cross validation by calculating the Adjusted Rand Index, classification accuracy and the macro F1 score at the single-cell level (Methods, Figure 3A). We left out one study at a time and clustered the cells from the other 18 datasets using marker sets and inferred labels of the held-out study based on their clustering near the closest centroid. We next calculated the ARI value for each study this way. From Figure 3A, we observe that overall, calculated marker sets demonstrate a higher single-cell LOO ARI value, than the canonical marker set across 9 out of the 19 datasets (Supplementary Figure S6). The average ARI score for the calculated set is 0.32 ± 0.11 S.D., comparable to that for the canonical marker set (0.34 ± 0.08 S.D.) (Supplementary Table S5). Since clustering was performed with only the calculated markers, these results suggest that the calculated markers can generalize better to different parts of the human cortex (VIS, dlPFC, PFCV, PFCA) compared to canonical markers and demonstrate nearly equal performance in some regions (CgG, PFCK, S1lm, M1ul, MFC). Canonical markers seem to outperform the calculated markers in regions like ACCK, A1C and S1ul.

**Figure 3:**
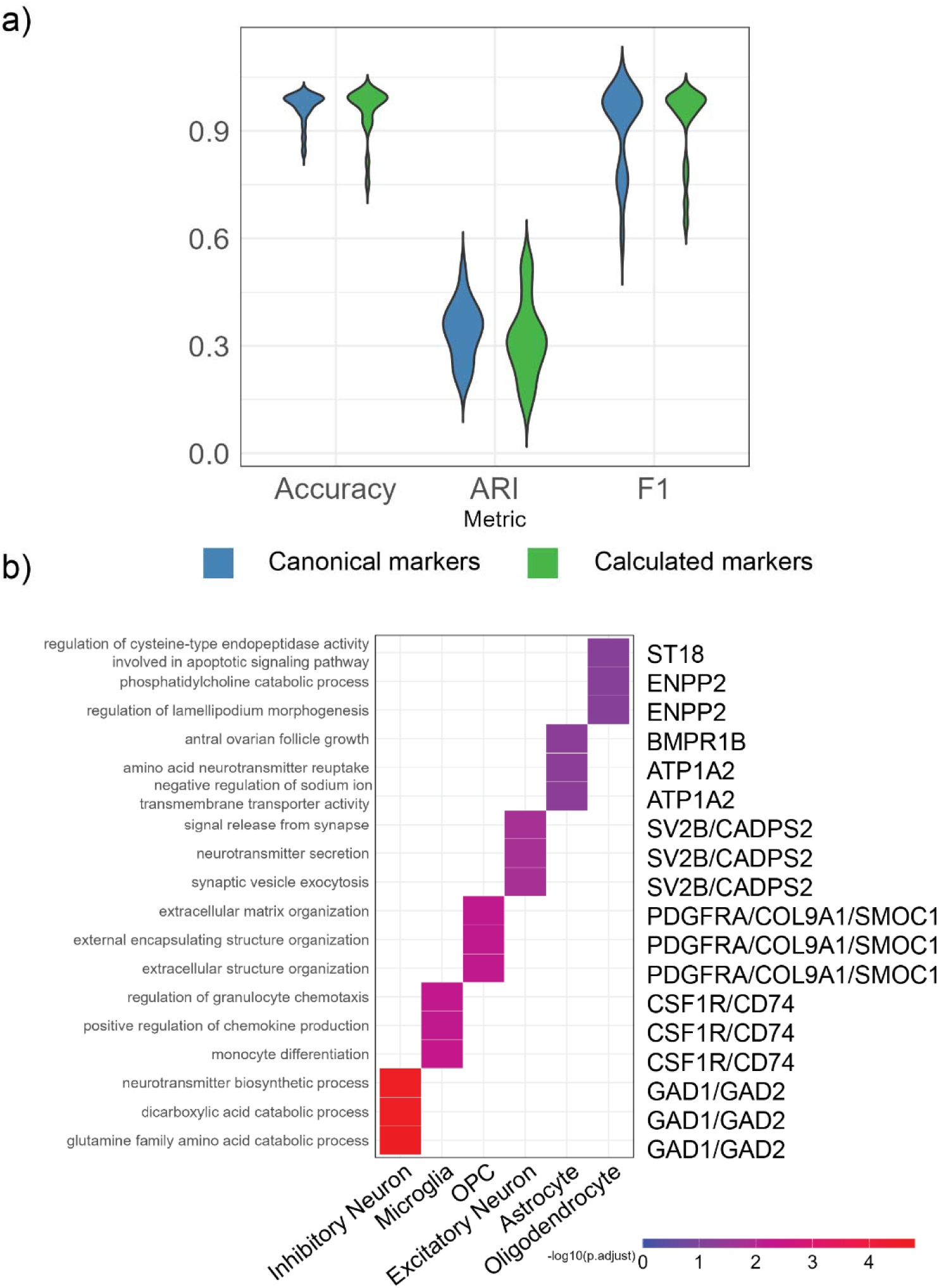
A) Calculated marker sets are generalized for clustering and classification tasks and enrich for relevant BP terms with canonical markers. Calculated marker sets demonstrate comparable ARI, accuracy and averaged macro F1 score as canonical markers. B) Top Biological Process Gene Ontology terms enriched using the Calculated marker sets vs the universe of 14118 genes (adjusted p-value < 0.05).

We further validated the classification performances of calculated and canonical marker sets. Since we are performing multi-class classification and there is an imbalance between the number of cells in each class, we used the F1 score, the harmonic mean of the precision and recall, to assess the classification performance in a balanced way by considering both false positives and false negatives using precision and recall in its calculation. We calculated the F1 score for each cluster and averaged it to obtain the average macro F1 score. We trained a multinomial logistic regression classifier using *glmnet()* and calculated accuracy and macro F1 scores (Methods) using the predicted labels of the held-out set (Figure 3A). Even though both marker sets have good classification performance accuracy (Figure 3A), calculated markers offer a higher clustering score than canonical markers in 15/19 datasets and a higher averaged macro F1 score in 12/19 datasets (Supplementary Figures S6-8). We further performed Gene Ontology Enrichment analysis to determine biological processes enriched by the calculated marker set. BP terms associated with chemokine production and response to external stimulus were enriched by microglial markers, while oligodendrocyte markers enriched for migration and lamellipodium morphogenesis (Figure 3B, Supplementary Table S6).

### High fidelity regional neuronal subtype level markers

Motivated by the success of our approach in expanding the list of canonical cell type markers for the 6 major cell types, we attempted to extend the analysis to cortical cell subtypes. Among all the studies considered in our meta-analysis, almost all studies annotated cells at the subtype level. Layer specific neuronal subtypes for both excitatory and inhibitory neurons were the most common subtype annotations available, while a few glial cell subtypes have also been addressed in some studies. Since cell subtype nomenclature was not consistent across studies, we decided to provide region-specific estimates of cell type marker lists. Consequently, we have used only the local components of the above methods, namely - *Adjusted p-value, log2FoldChange, fidelity* and *background.* For each cell type, we selected genes with *Adjustedp-value < 0.05, log2FoldChange > 1, fidelity > 80%* and *background < 15%.* We then prioritized fidelity and selected the top 15 genes with the highest fidelity from this list.

After performing differential expression analysis for each available subtype in all studies, we fine-tuned the results to obtain high fidelity subtype markers for each region (Methods). We then calculated the average expression fidelity and background using the corresponding metrics from all cell subtypes in each region (Figure 4A, B, Supplementary Figures S9-12). We provide the marker genes used to achieve this fidelity and background levels as calculated layer specific markers from each region (Supplementary Table S7).

**Figure 4:**
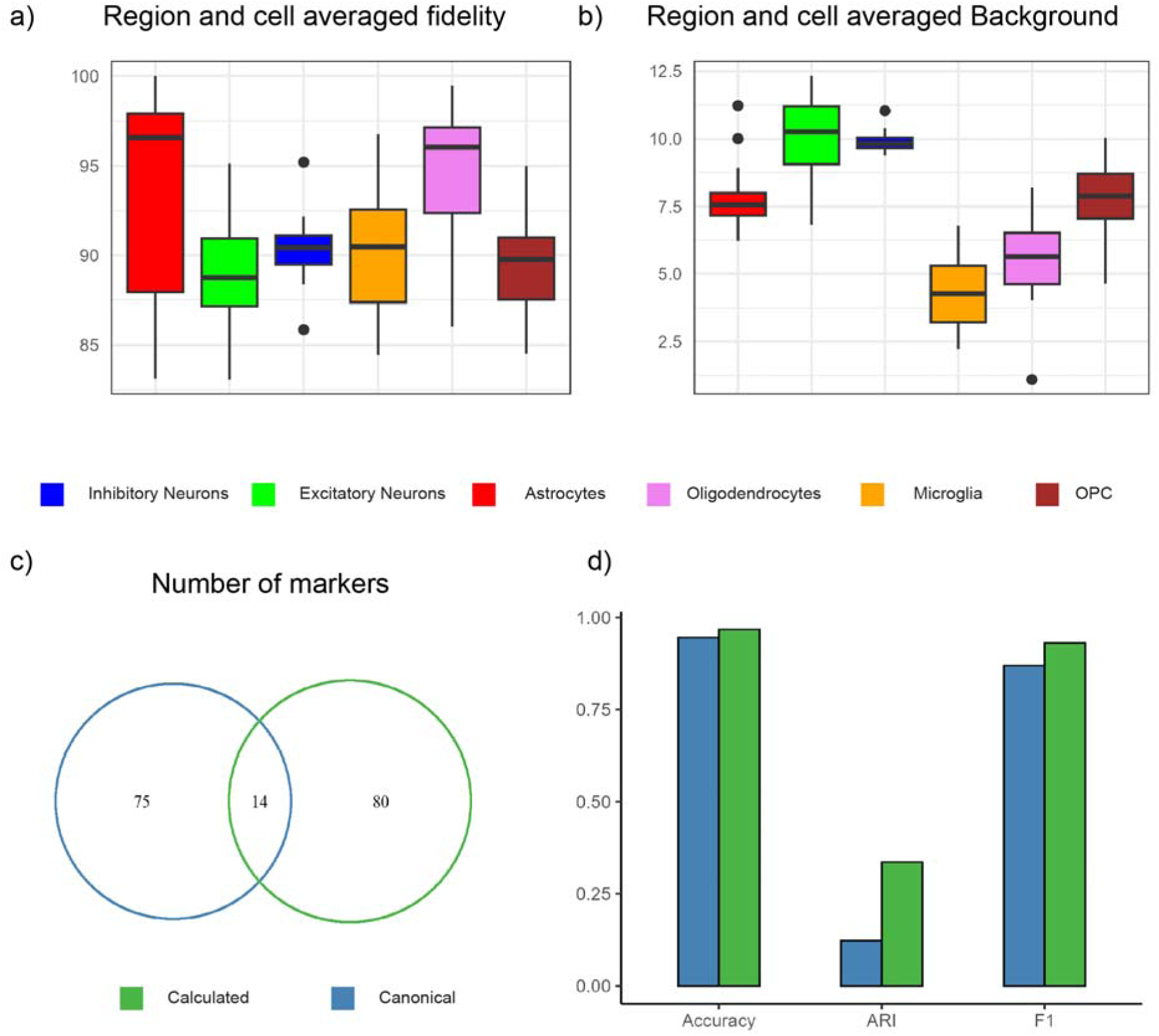
Calculated region-specific subtype markers demonstrate high fidelity and low background. A) Average fidelity and B) background percentages of all calculated subtype markers of the 6 major cell types in all regions B) Venn diagram depicting the number of new subtype markers identified from this analysis, with 14 markers overlapping with the original set. D) Accuracy, ARI and F1 metrics from the validation step calculated in both clustering and classification of cells from the dlPFC dataset.

To validate these subtype markers, we selected the dataset from the dorsolateral Prefrontal Cortex, which has a high number of cells and author provided markers (Ma et al.). Using the provided metadata, we chose 89 total layer specific markers. In comparison to this set, we selected the top 15 markers for each cell subtype from our method mentioned above for a total of 94 proposed markers (Figure 4C), of which 75 were different from the original 89 marker genes. We validated the marker sets (Methods) by calculating the accuracy, ARI, and F1 macro scores (Figure 4D). We observe that fidelity and background informed layer-specific markers outperformed the canonically used subtype markers in all three metrics. These results indicate that a fidelity-based statistical approach such as ours has the potential to improve our understanding of layer-specific markers in the human cortex.

## Discussion

Canonical cell type markers have been foundational to illuminating cell type specific molecular features of the human brain. Advances in single cell resolution studies have expanded our understanding of cell type heterogeneity in the human brain and established a combinatorial perspective of gene expression programs of cell types. Nevertheless, canonical cell type makers are critically employed in the annotation of cell type clusters in single-cell expression data, as well as in targeted validation studies. In this study, utilizing the information from cellular resolution studies, we investigated the variability of canonical cell type marker expression indifferent regions of the human brain. We first focused on the six major cell types, given that it is relatively straightforward to identify them in different research contexts, thus providing an opportunity to examine how variable is gene expression and how we can extend the concepts of cell type markers in the single cell era.

As expected, gene expression levels are highly variable in different cortical regions. In addition, we demonstrate that marker gene expression in the specific cell type (which we termed as the fidelity) as well as expression in unrelated cell types (which we termed as the background expression) vary substantially from one part of the cortex to another. (Figure 1B). While we confirmed that all canonical markers are significantly differentially expressed in the corresponding cell type cluster (*adjusted p-value < 0.05* and *log2FoldChange >> 1*), there is a large variation in gene expression fidelity. For instance, we noticed that the *GFAP* gene, which is a commonly used immuno-staining marker for astrocytes, was expressed in less than 50% of astrocyte cells in 7 cortical regions, while the solute carrier family member-based astrocyte markers *SLC17A7* and *SLC4A4*, had high (>90%) fidelity in 13 regions. However, these genes were also expressed in many cells that are not in the astrocyte cluster, but in other clusters. In other words, they were subject to significant background levels. The *AQP4* marker also had low fidelity levels in the Posterior Cingulate Cortex (PCC) and the Visual Cortex (VIS). The *MOG* oligodendrocyte markers also had reduced fidelity levels in the PCC and VIS regions. The fidelity levels of the microglial markers *AIF1* (7.16%) and *CSF1R (67.9%)* were reduced in the dlPFC dataset relative to the remaining considered datasets. Similarly, the microglial markers *APBB1IP* had a low fidelity level (59 %) in the visual cortex (VIS). The *PDGFRA* marker used to distinguish OPCs also showed lower fidelity levels in dlPFC, PCC and one of the Prefrontal Cortex (PFCV, < 50%) datasets. As different parts of the human cortex are specialized to perform specific tasks, we anticipate that gene expression programs, including the regulation of canonical marker gene expression and proportions of cells expressing these genes, would be different.

Another reason for this is that since canonical markers are highly expressed genes, and highly expressed genes in general tend to be highly expressed in other parts of the cortex as well. In other words, the correlations between expression and background tend to be positive. And in some cases, genes with high fidelity also have high background expression (Figure 1C). However, Figure 1C also indicates that some genes are highly expressed and exhibit high fidelity, without strong background expression. We sought to identify and expand the list of cell type markers that may be used to distinguish major cell types. Recent benchmarking studies have shown the effectiveness of differential expression testing over various feature selection methods ^24^ in single-cell marker detection methods. In this study, we use a combination of pseudo-bulk differential expression testing and the resolution offered by single-cell expression fidelity to discover highly specific markers for the six major brain cell types in the human cortex.

In other words, we introduce the idea of complementing marker discovery and feature selection methods by explicitly utilizing the fidelity and background expression. As gene expression is highly variable across different cortical regions and cell types, to discover novel biomarkers, thresholds of fidelity, background and Cohen’s d-metric had to be tuned in a cell-type specific way, further emphasizing the highly heterogeneous nature of cell types and their expression landscapes in the human cortex. Our method re-discovered both inhibitory neuronal markers - *GAD1* and *GAD2* and 6 other canonical markers. This suggests that the novel markers we discovered might also be biologically relevant. Consistently, Gene Ontology analysis yields enriched BP terms containing combinations of both calculated and canonical markers.

We utilized various performance evaluation metrics to rigorously validate the performance of the calculated markers, including Leave-One-Out methods for clustering and generalizability across different cortical regions and classification, demonstrating the improvement using concepts of True Positive Rates and False Positive Rates for both sets of markers (Supplementary Figure S6-S8). We observed that our calculated markers generalize better than or comparable to canonical markers in more than 9 different cases out of the total 19 datasets, while preserving comparable performance in classification at the single-cell and pseudo-bulk level. However, there are still regions where the clustering performance needs improvement, such the frontal cortex. It is also important to note that canonical cell type markers show similarly low performance in these regions. Additional studies are needed to understand regional effects influencing marker performance.

We further provide access to our methodology to the community to discover these markers, test them, and explore other sets of markers which might be better suited to their requirements. The interactive application we developed also summarizes the performance of genes of interest in different cortical regions (Tab: Gene region ranking) in the application interface). We also provide a feature plot option to visualize pseudo-bulk expression levels of desired genes. This will enable the scientific community to select multiple markers for histological staining and fluorescent separation using antibodies in an evidence-based manner factoring in fidelity and background levels in a region-specific manner.

As single-cell technologies grow in resolution of molecular profiling, there is a growing interest in annotating cellular subtypes. The cerebral cortex being a highly heterogeneous tissue displays a large family of neuronal subtypes and layer-specific neuronal markers are not easy to define and validate. By following a fidelity-based approach, we showed that we can discover novel subtype markers that could complement our current understanding of layer-specific neuronal marker genes and improve upon clustering and classification tasks. Further analyses of these subtype markers may reveal additional insights into understanding gene expression programs that constitute the complex human brain layer-specific cell types. We propose that fidelity-based approaches such as our method become increasingly relevant in histological separation and isolation of cells, particularly since the number of layer-specific cells extracted in each experiment is low.

## Methods

### Data Source

We collected published single-cell and single-nuclei RNASeq raw count matrices from several published studies spanning various regions of the human cortex. We performed this meta-analysis using a total of 12 different studies (Bakken et al.; Brenner et al.; Lake et al.; Khrameeva et al.; Velmeshev et al.; Yang et al.; Jorstad et al.; Caglayan et al.; Kanton et al.; Ma et al.) and the multiple cortical regions dataset from the Allen Institute for Brain Science (*Dataset: Allen Institute for Brain Science (2021). Allen Cell Types Database -- Human Multiple Cortical Areas [Dataset]. Available from* Celltypes.Brain-Map.Org/Rnaseq) spanning a total of 16 unique cortical regions and a total of 19 datasets as some studies addressed the same region. Supplementary Table S1 details the full list of these studies, the size of the dataset and the associated study. We retrieved the data from the provided universal resource locator links.

### Single cell processing

Using the data in the originally published state, we first used CellBender (Fleming et al.) to remove ambient RNA contamination. Due to the large size of some matrices, we reduced the memory requirement for this step by selecting only the protein coding genes according to GRCh38. We performed this for the datasets addressing the M1 cortex, MTG, Multiple Cortical Regions (after splitting the matrix into constituent regions), We converted these matrices into the .HDF5 format using the R function - write10xCounts(). We also split the datasets MTG (Jorstad et al.) and into constituent samples and selected only protein coding genes to reduce computation times. We performed ambient RNA removal using the default parameters of the CellBender command and specified approximate estimates for the number of expected cells (approximately 80% of total number of cells) and included droplets based on the sample size. The CellBender processed human datasets from our earlier studies on the datasets for PCC (Caglayan et al.), dlPFC (Ma et al.) and PFC (Kanton et al.) were re-processed using the updated version of CellBender to maintain consistency with the above datasets.

We next performed single-cell quality control to remove cells expressing more than 5% mitochondrial DNA, cells with less than 200 genes detected (*nFeature_RNA*) and genes expressed in less than 3 barcodes from each dataset. We further removed doublets via automatic doublet finding using DoubletFinderV3() (McGinnis et al.) which has been modified for Seurat version V5 (URL: https://github.com/chris-mcginnis-ucsf/DoubletFinder/issues/161source?), from the DropletUtil library. (See Supplementary Table S1 for these studies and Supplementary Table S2 for the final number of barcodes from each region).

We pseudo-bulk aggregated all barcoded cells from the remaining studies by summing the raw transcript counts grouped by cell type and sample based on the cell type annotations provided by the authors in their respective studies. We aggregated them by the six traditional brain cell types; Astrocytes, Excitatory Neurons, Inhibitory Neurons, Microglia, Oligodendrocytes, OPCs.

We next generated pseudo-bulk clusters for one-vs-all differential expression analysis where we took out one cell type at a time and pseudo-bulk aggregated by summing all remaining pseudo-bulk clusters. We performed differential expression analysis using DESeq2 (Love et al.) after normalizing the counts using the median of ratios technique, followed by the Wald Test and successively performed multiple testing correction using the Benjamini-Hochberg method (Benjamini and Hochberg) of adjusting the resulting p-values.

### Pseudo-bulk DE marker list generation framework

#### Global Cohen distance calculation

The global Cohen distance is a metric used to evaluate the distance between the means of the distribution of log-transformed pseudo-bulk expression levels of a given marker gene associated with a cell type of interest vs the mean of the distribution of log-transformed pseudo-bulk expression levels of the background (cell types other than the current one) (Cohen). To calculate the Cohen distance for a given marker gene, we first calculate the pooled standard deviation:

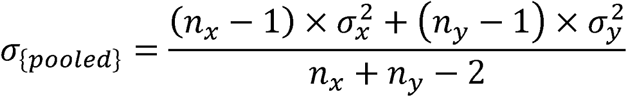

Where nx and ny are the number of samples in the two distributions and 2x and 2y are the variances of the two distributions. We use the pooled standard deviations to calculate the Cohen-d distance:

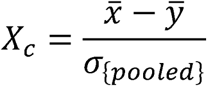

Where x and y are the means of the two distributions.

To extend the considered list of canonical cell markers, we developed a statistical framework considering both local and global performance metrics, leveraging both the single-cell and pseudo-bulk resolution. Motivated by extensive studies conducted in benchmarking marker discovery methods (Pullin and McCarthy), we first performed one-vs-all differential expression analysis (DESeq2) to determine cell type specific gene lists from each data set and applied several criteria to better identify cell-type specifically expressed genes. From the list of tables of differentially expressed genes, we begin filtering by selecting genes which were significantly differentially expressed (adjusted p-value < 0.05) and *log2FoldChange > 2* and applied cell-specific fidelity and background thresholds (Methods). We then select the top 5 most frequently occurring genes, meeting the above criteria. We then selected genes using cell-type specific Cohen distance thresholds. The final values for these cell-specific parameters were determined using a trial-and-error approach and have been tabulated below:

**Table 1.** Cell type specific thresholds used for marker selection for each of the 6 major cell types.

|  | Fidelity threshold | Background threshold | Cohen distance |
| --- | --- | --- | --- |
| Astrocyte | 80 | 15 | 2.25 |
| Excitatory Neuron | 80 | 15 | 0 |
| Inhibitory Neuron | 90 | 5 | 2 |
| Microglia | 95 | 5 | 4.5 |
| Oligodendrocytes | 95 | 10 | 1 |
| OPC | 90 | 5 | 2 |

### Marker validation methods/generalizability

#### Single-cell Leave-One-Out clustering cross validation

To demonstrate the generalizability in clustering by the calculated cell markers across multiple regions, we performed Leave-One-Out cross validation. In this method, for each fold, we use the cells from one dataset as a hold-out and perform clustering on all the cells of the other 18 datasets. Here, we normalize the full dataset by scaling it by 10,000 and by the library size (sum of all cells) and then apply a log transformation. After clustering in this transformed space (using Louvain clustering), we grouped the cells in the hold-out set to the centroid nearest to it cand assigned labels to them based on this grouping. We then compare the assigned labels to the true labels and calculate the adjusted rand index (ARI) using the *adjustedRandIndex()* function from the *mclust* library. This conveys how well the markers can cluster cells from new cortical regions. We performed the clustering for 10 different seeds chosen at random and calculated the average ARI score using all seeds. The confidence intervals in Supplementary Figure S6 were calculated as following standard methods (Casella and Berger):

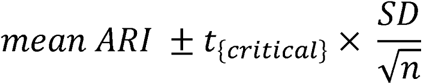

For subtype level clustering, we used the top 20 principal components and the hdbscan clustering algorithm for faster computation since gene sets were larger and clustering was computationally expensive. We used a 70-30 train-test split.

#### Single-cell Leave-One-Out classification cross validation

To investigate whether markers classify cells correctly, we performed a similar Leave-One-Out classification cross-validation. In this approach, we used the same single-cell normalization as above and the calculated and canonical marker sets to train a regularized multi-logistic regression classifier using *glment* (Friedman et al.; Simon et al.). We trained the classifier on all regions barring the hold-out set and then used the trained model to predict the labels of the cells in the hold-out set. We used a separate seed for each hold out set, but maintained the same seed for comparison between calculated and canonical marker sets. We used the classifier to calculate classification accuracy and detected the true positives, false positives and false negatives to calculate the mean macro-F1 score. We used the conventional definition of precision and recall as ((Powers; Sokolova and Lapalme)):

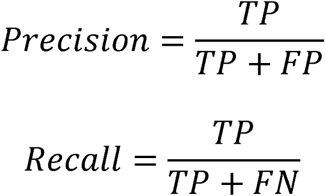

We then calculate the macro-F1 score for each cell type as:

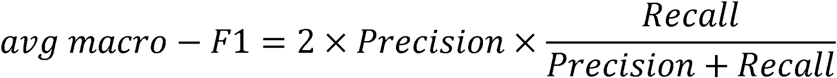

We then calculate the mean of this metric across all 6 cell types and report this as the mean macro-F1 score.

### Pseudo-bulk marker ARI and silhouette scores

We used three metrics to evaluate the performance of calculated marker genes from the above pipeline at the pseudo-bulk level, we used two metrics - the adjusted rand index, the silhouette value

#### Adjusted Rand Index

We first downsample the pseudo-bulk aggregated count matrix to consider only the specific genes of interest. We scaled the data and calculated the distance matrix using the euclidean distance and clustered the samples using the ward.D2 method. We next predicted the labels using the *cutree* function, using this dendrogram and using 6 clusters for the k parameter. We then used the original matrix columns and the predicted labels to calculate the Adjusted Rand Index (ARI) (Rand) using the *adjustedRandIndex()*.

We selected gene lists that had a comparable or better ARI value to that of the canonical marker superset. The ARI score is an external clustering score that conveys both the clustering capacity and the classification accuracy of the calculated marker set.

#### The Average Silhouette Score

We calculate the average silhouette score (Rousseeuw) using the predicted labels and the distance matrix used in the ARI section in the *silhouette()* function in R. The silhouette score indicates how similar a sample is to its own cluster (cohesiveness) and how different it is from other clusters. We calculated the silhouette value of each sample in the dendrogram and calculated the average silhouette score for each calculated marker set. We took this metric into account and selected calculated marker gene sets with comparable silhouette scores to the canonical marker set.

### Gene-Ontology enrichment

All Gene Ontology (GO) enrichment, both for the cell type markers, were performed using the enrichGO() function from the ClusterProfiler library R (Yu; Wu et al.), using the 14,118 common genes detected in all studies as the background, specifically focusing on Biological Processes. We used *Benjamini-Hochberg* multiple testing correction, with a *pvalueCutoff < 0.05* and *qvalueCutoff < 0.05*.

### Subtype marker list generation

To discover marker genes for cell subtypes, we first performed differential expression analysis at the subtype level from all studies. In all, 17 studies provided subtype resolution at various levels. Since the nomenclature of cell subtypes was not consistent across all studies, we performed a local analysis for each study. Similar to the major cell types, we pseudo-bulk aggregated by cell subtype and by sample and used this data to calculate tau-specificity, fidelity and background levels. We then removed samples with less than 40 cells to arrive at a global matrix of 797 samples. We then selected genes with p-adj < 0.05, background < 15%, fidelity > 80 %, and log2FoldChange > 2. We then further sorted by decreasing fidelity and selected the top 20 genes in the resulting table. We further averaged the resulting fidelity and background of these gene lists from each subtype grouped by major cell type from each region and used the resulting values to prepare Figure 4B, C and Supplementary Figures S8-11.

### Data Availability

The full single-cell cortical count matrix (488,397 cells) post QC, and associated cell metadata compiled from all 19 datasets is a great resource to build and test cell classification algorithms and is made publicly available at: 10.6084/m9.figshare.29992147. The pseudo-bulk aggregated count matrix prior to refinement is available for download from the GitHub repository of this project. Supplementary Figure S1 contains links of all source data prior to any QC, but intermediate datasets will be shared upon request - please directly contact. The RShiny application to assist in interactive biomarker discovery and testing is available at https://dmj6288.shinyapps.io/CortexMapperV1/. Supplementary Files (Figures and Tables) references in the paper are available on the Journal website and upon request.

### Code Availability

The software pipeline used to perform this analysis is publicly available at https://github.com/dmj6288/Expanding-canonical-cortical-cell-type-markers-in-the-era-of-singlecell-transcriptomics

## Acknowledgements

We thank Dr. Stephen Fleming (creator of CellBender) for detailed feedback regarding choice of CellBender parameters. We also thank all members of the Yi lab for discussions. We thank Karthik Somayaji, Shravan Muralidharan, and Sai Sukruth Bezugam (UC Santa Barbara) for lively conversations on machine learning and clustering.

## Author contributions statement

D.M.J. and S.V.Y. designed the study, conducted the analysis. D.M.J. developed the RShiny app. D.M.J. and S.V.Y. wrote the manuscript.

## Additional information

To include, in this order: **Accession codes** (where applicable); Not applicable

---

**Supplementary Figure S1:**
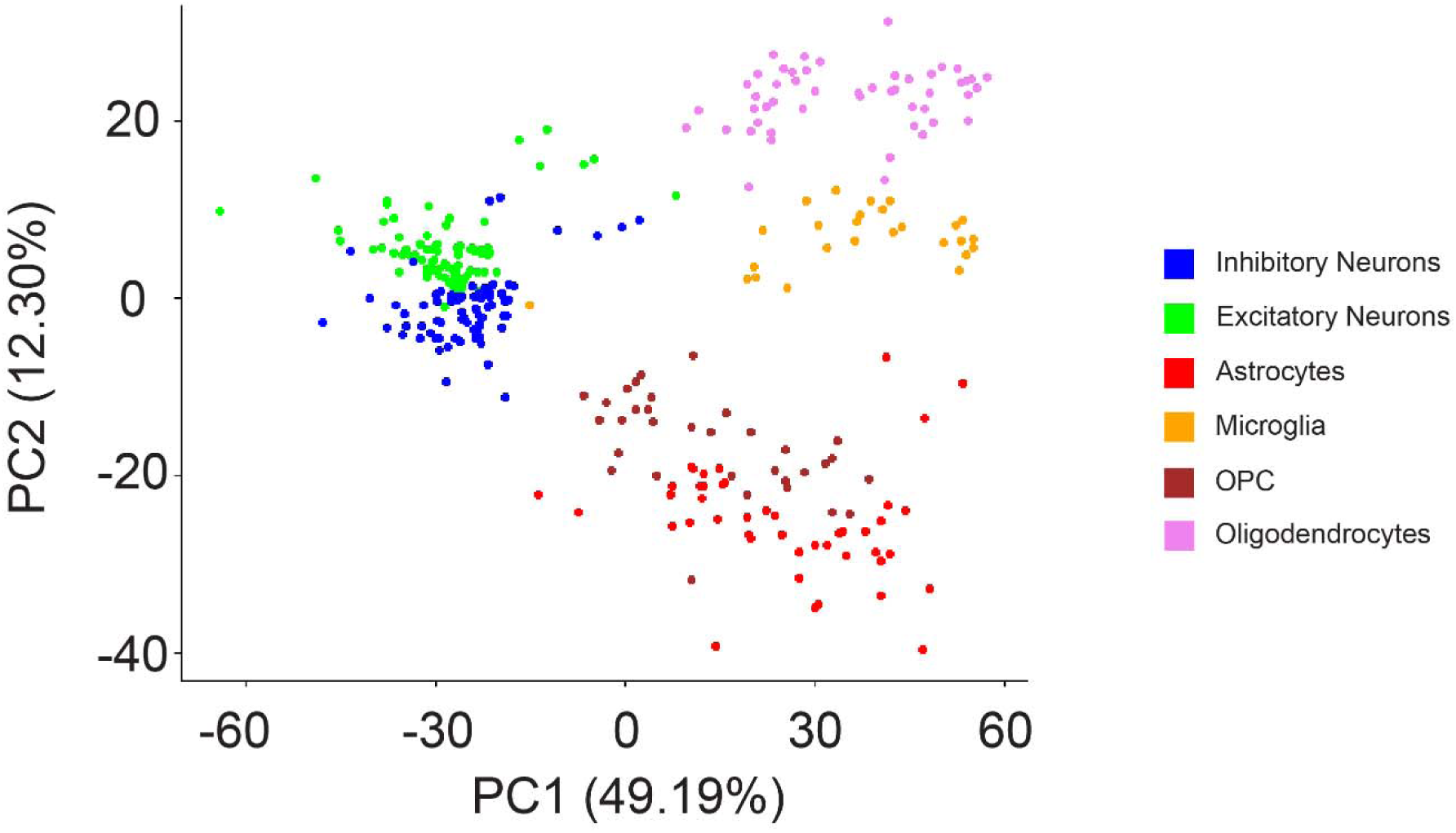
PCA plot of pseudo-bulk aggregated cell type-sample clusters show clear clustering in the first two PCs (calculated using DESeq2 normalized data, followed by VST transform and selecting the top 2000 variable genes), with both neuronal types in proximity, glial cell types close to each other and the microglial cluster apart from the 5 other cell types.

**Supplementary Figure S2:**
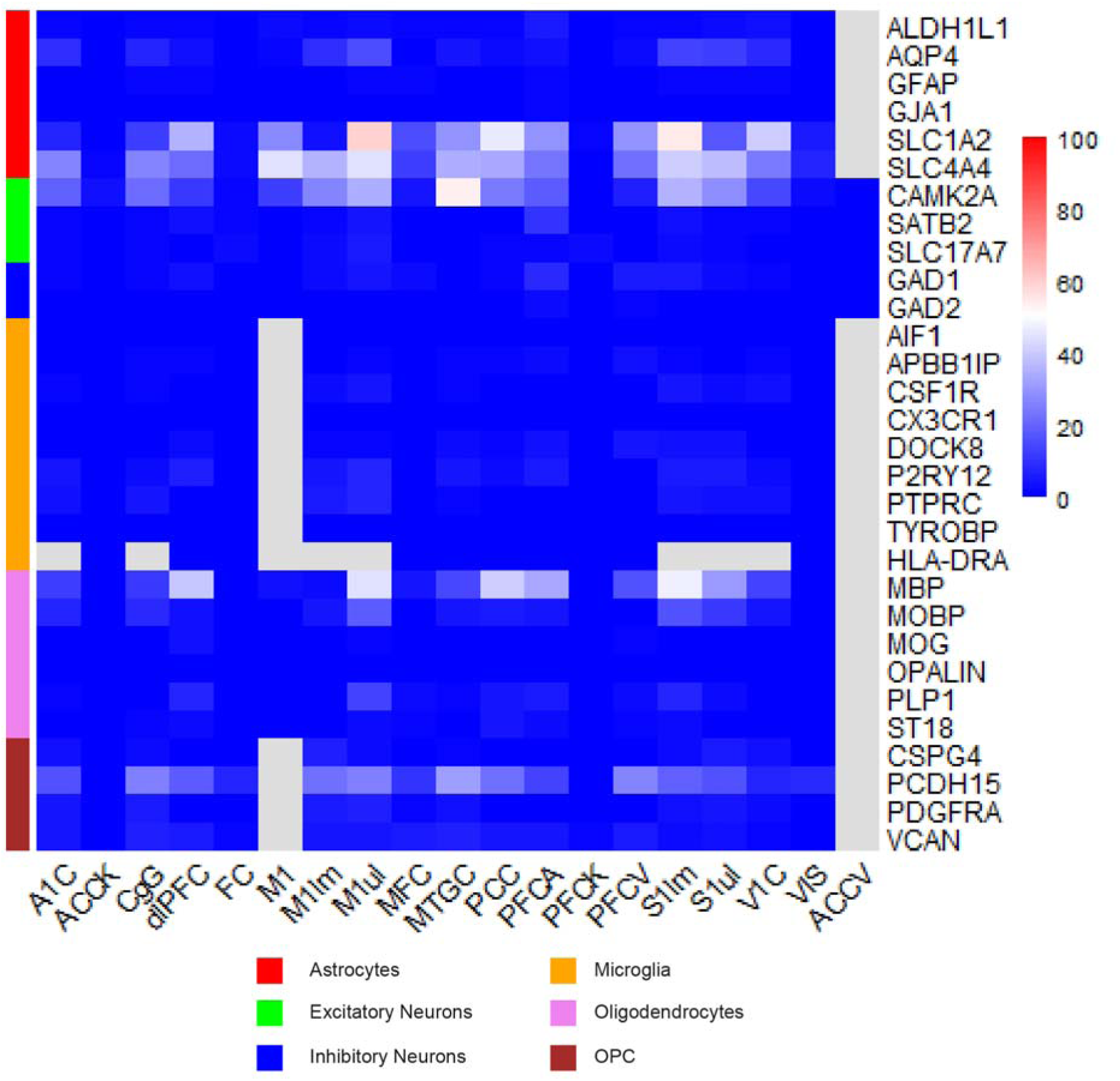
Some canonical cell type markers have significant background signals that vary across regions.

**Supplementary Figure S3:**
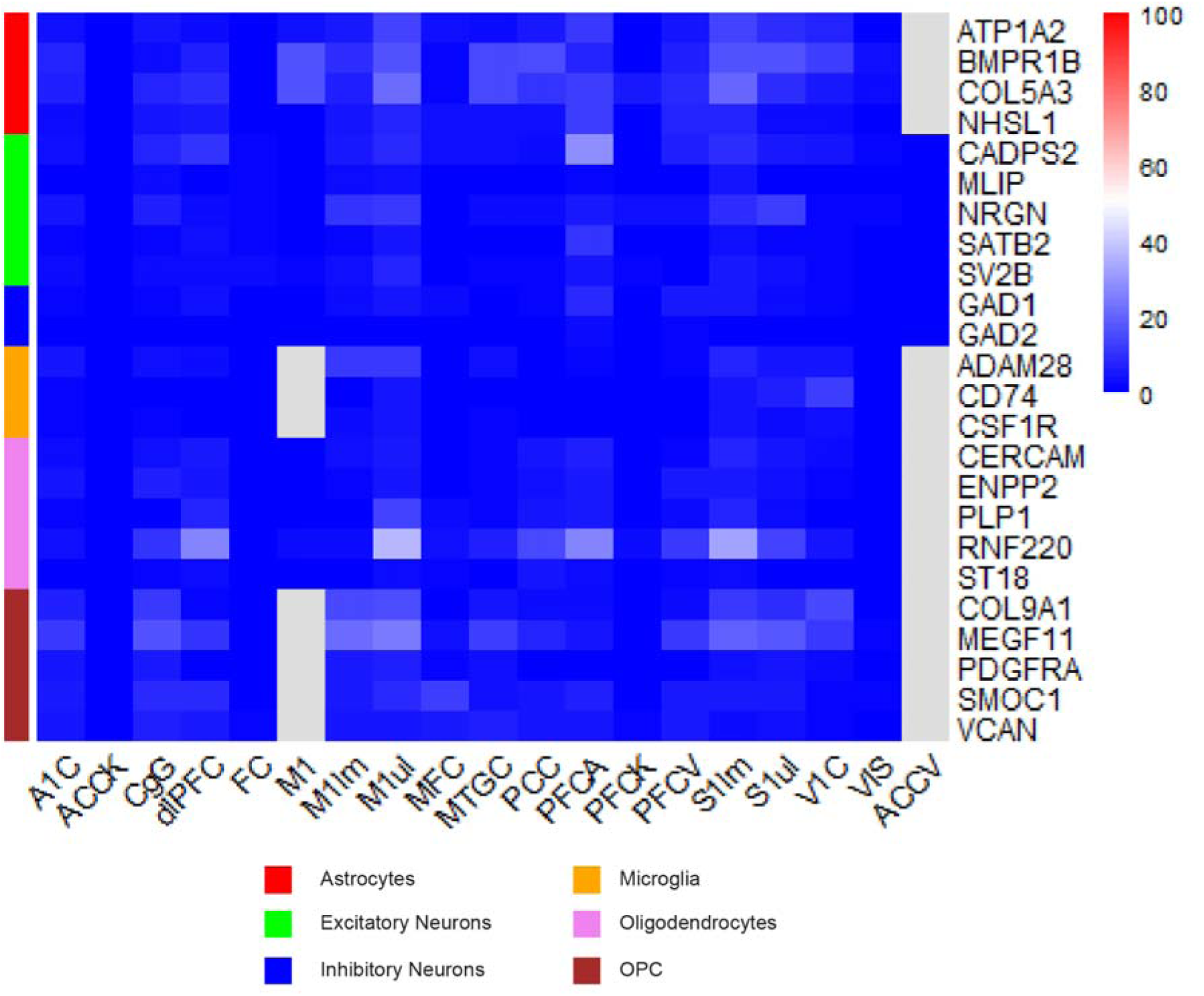
Discovered calculated markers have lower or similar background signals as canonical cell type markers.

**Supplementary Figure S4:**
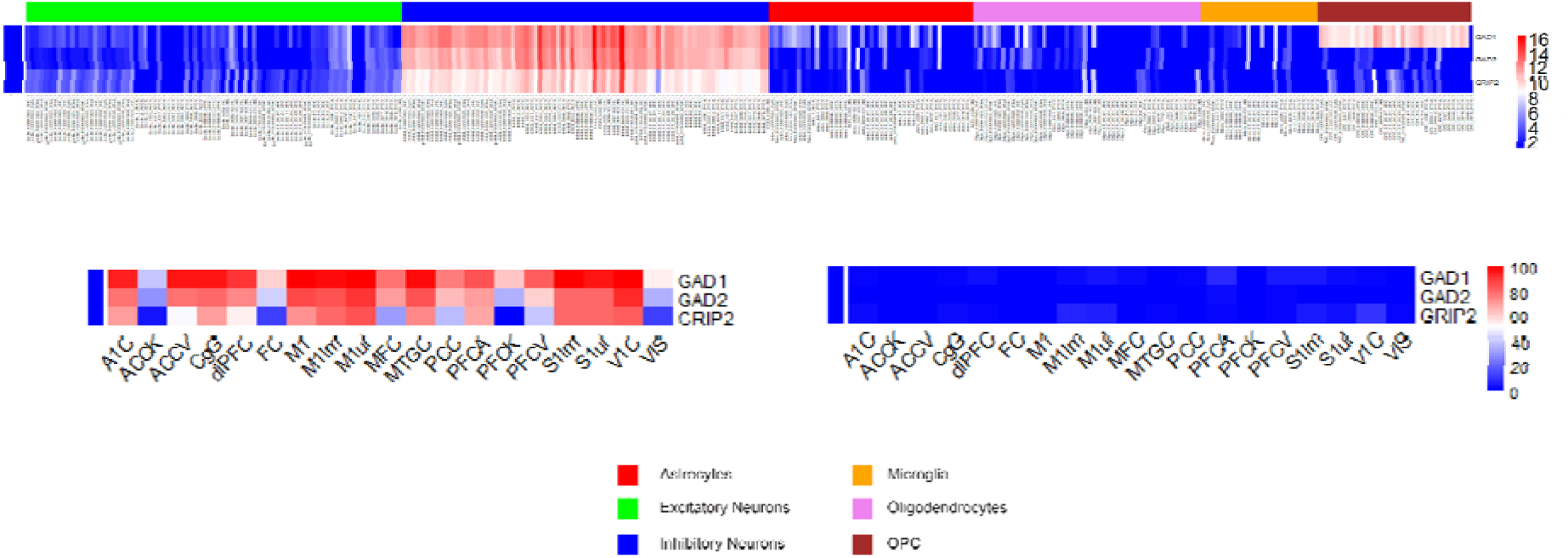
GRIP2 is a potential pan-inhibitory neuronal marker at par with GAD2, but has high variability in fidelity (lower left), but matches the background profile (lower right) of GAD1 and GAD2.

**Supplementary Figure S5:**
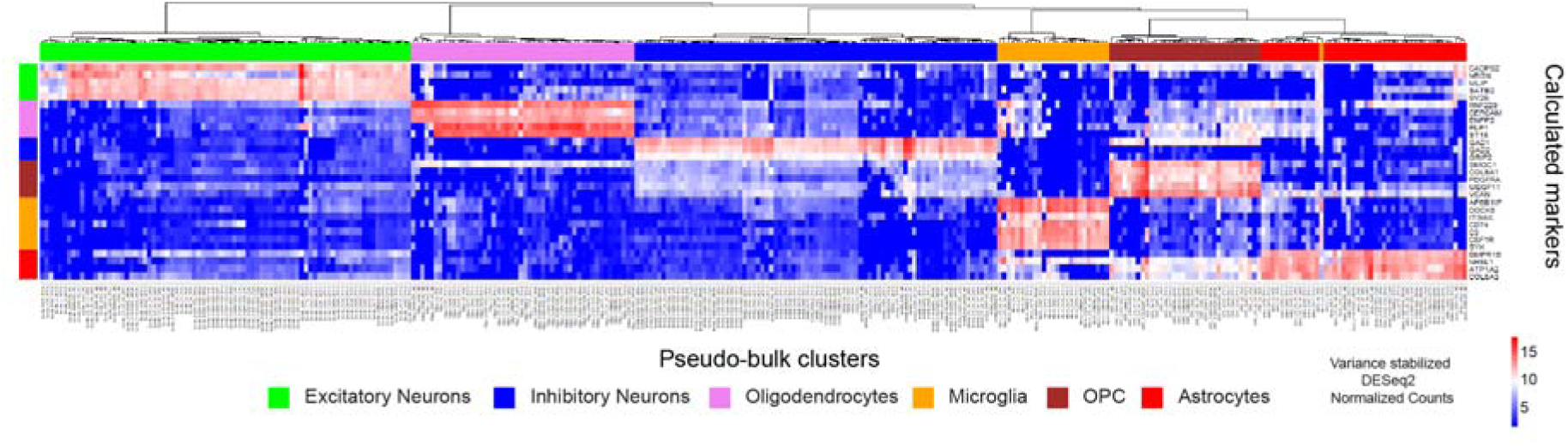
Calculated markers clustered pseudo-bulk clusters of cell types with high ARI and average Silhouette score.

**Supplementary Figure S6:**
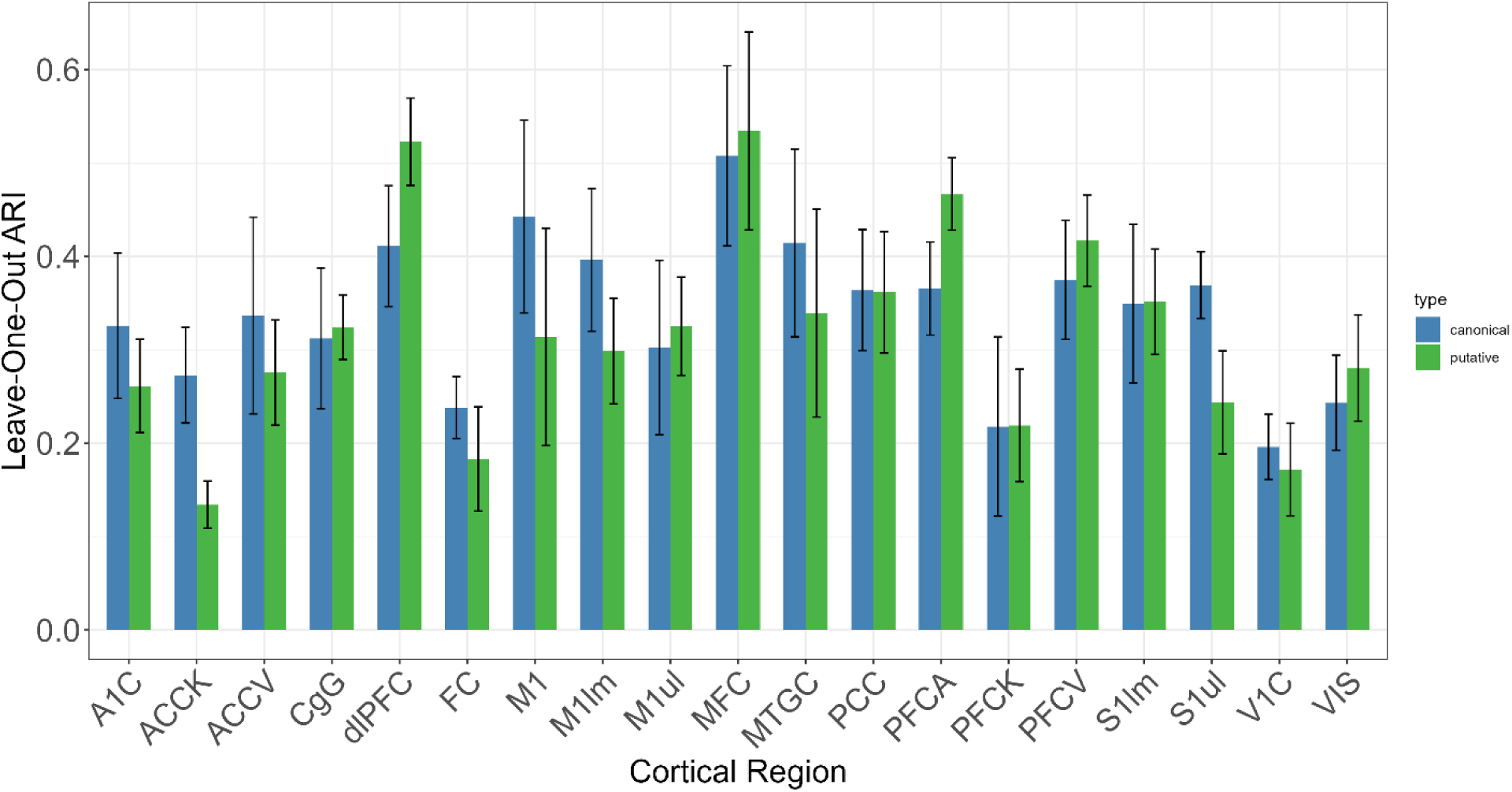
Calculated markers perform better or at par with canonical markers with higher ARI scores in 9/19 regions compared to canonical cell type markers.

**Supplementary Figure S7:**
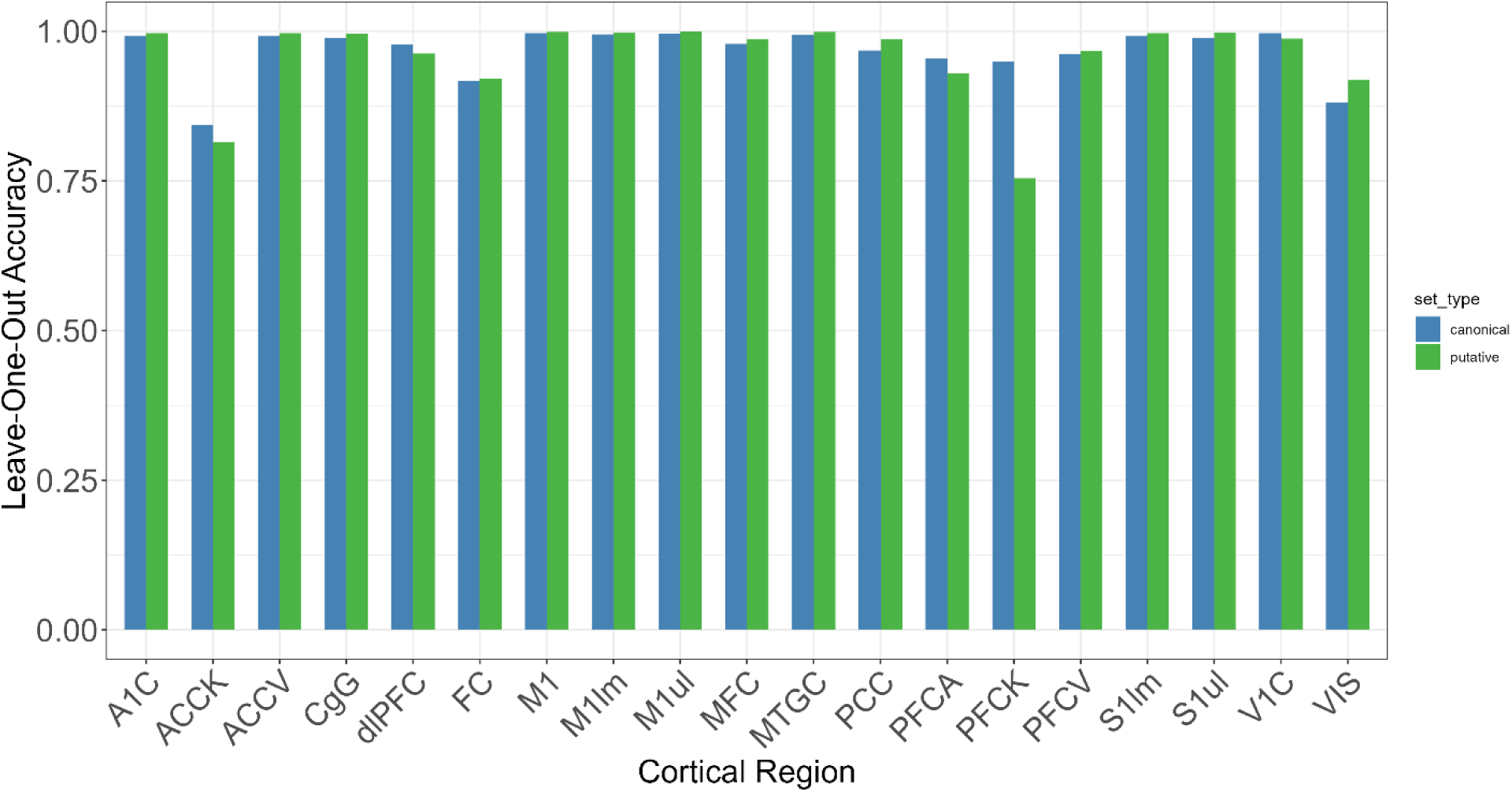
Calculated markers outperform canonical markers with higher accuracy scores in 15/19 regions compared to the canonical cell type markers

**Supplementary Figure S8:**
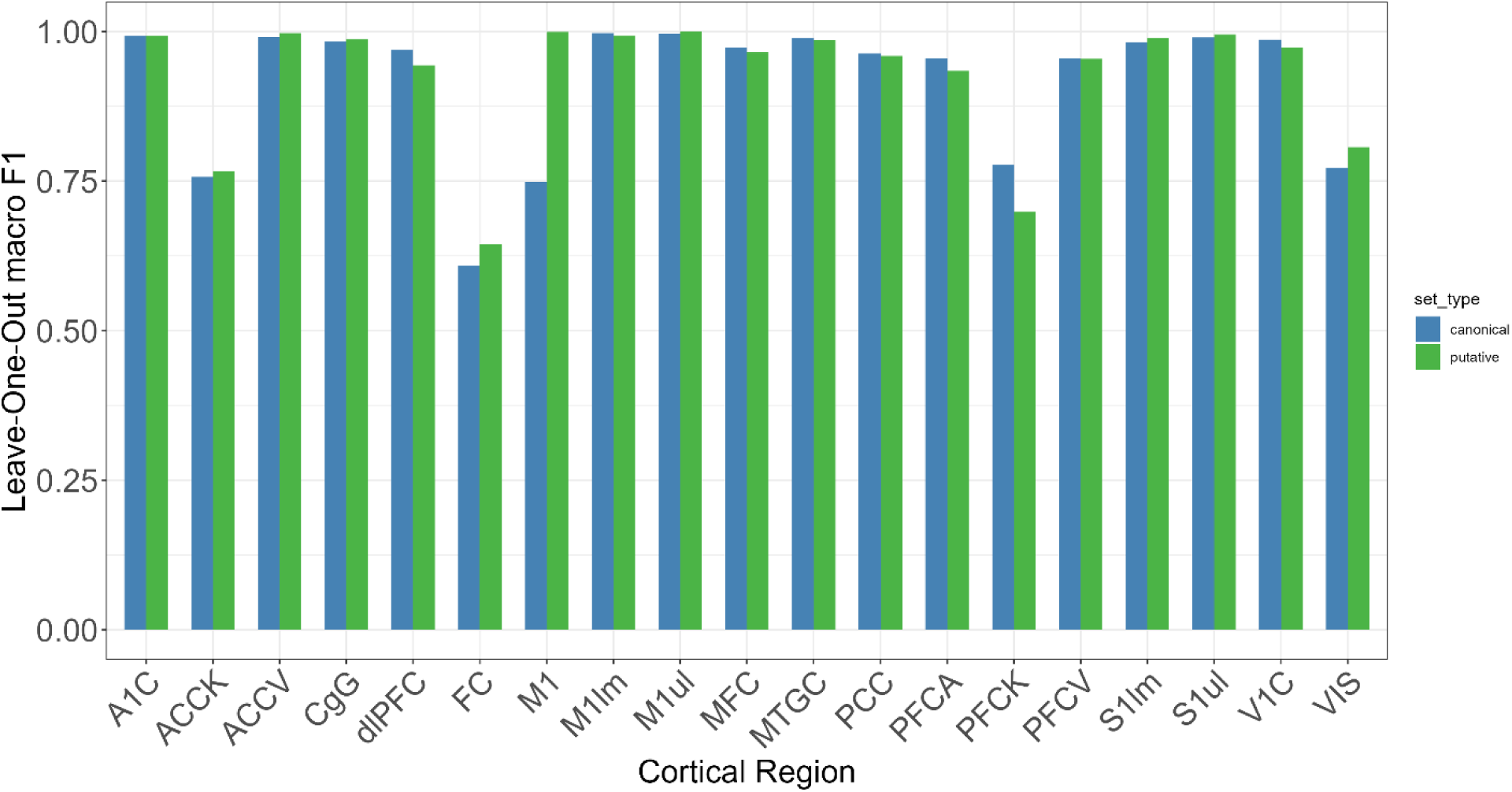
Calculated markers outperform canonical markers with higher average macro F1 scores in 12/19 regions compared to the canonical cell type markers

**Supplementary Figure S9:**
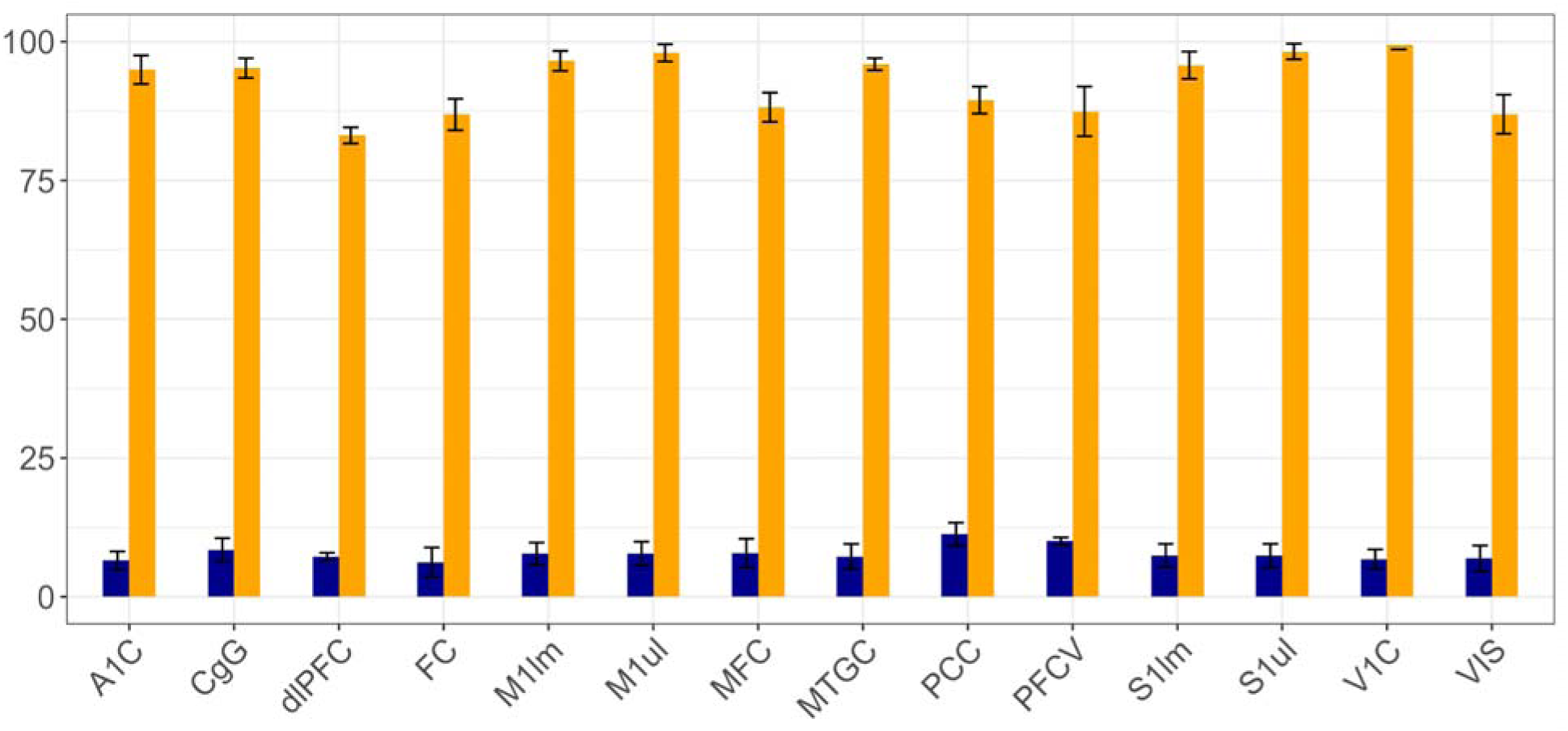
High fidelity and low background levels of astrocyte subtype markers calculated using our method across all studies with subtype resolution.

**Supplementary Figure S10:**
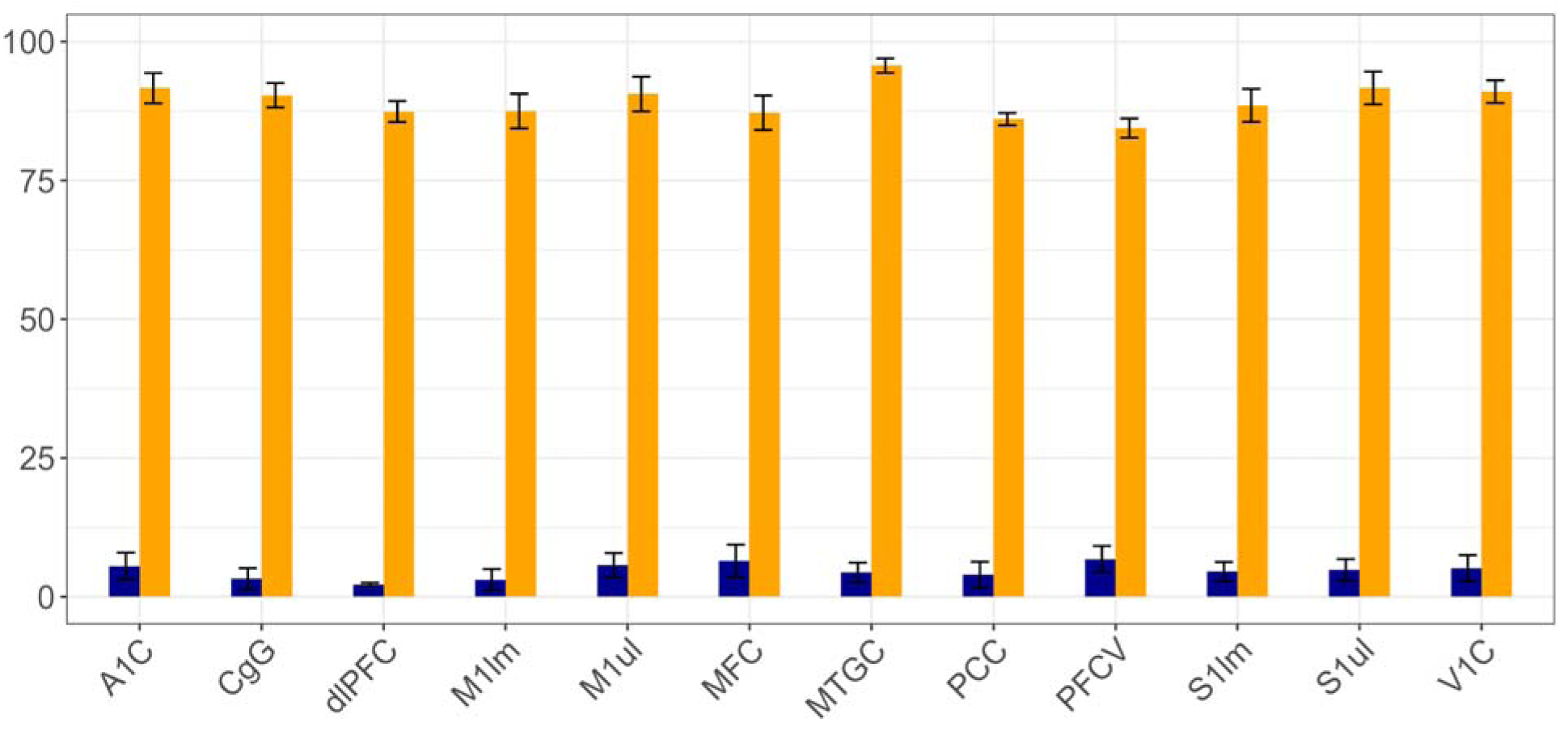
High fidelity and low background levels of microglia subtype markers calculated using our method across all studies with subtype resolution.

**Supplementary Figure S11:**
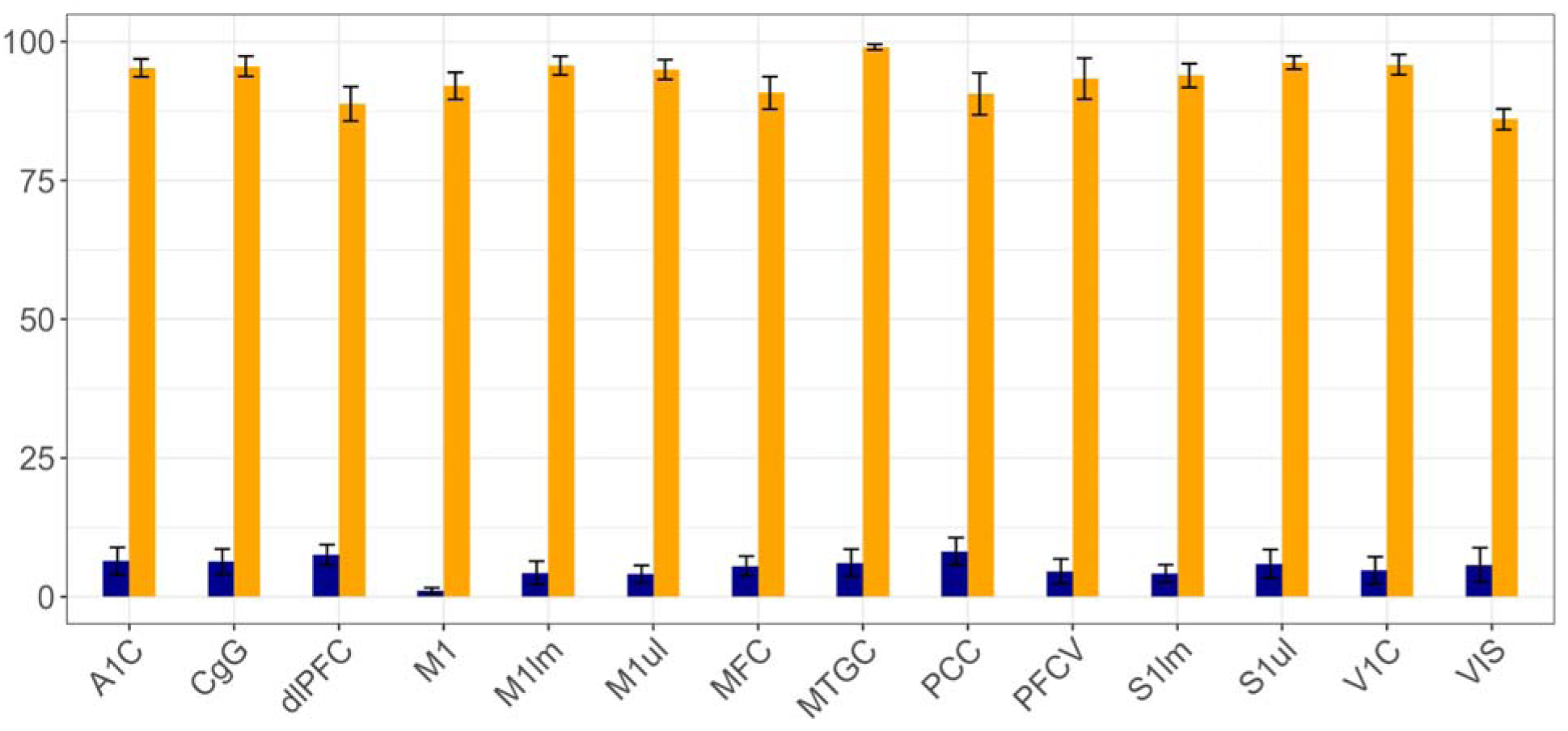
High fidelity and low background levels of oligodendrocyte subtype markers calculated using our method across all studies with subtype resolution.

**Supplementary Figure S12:**
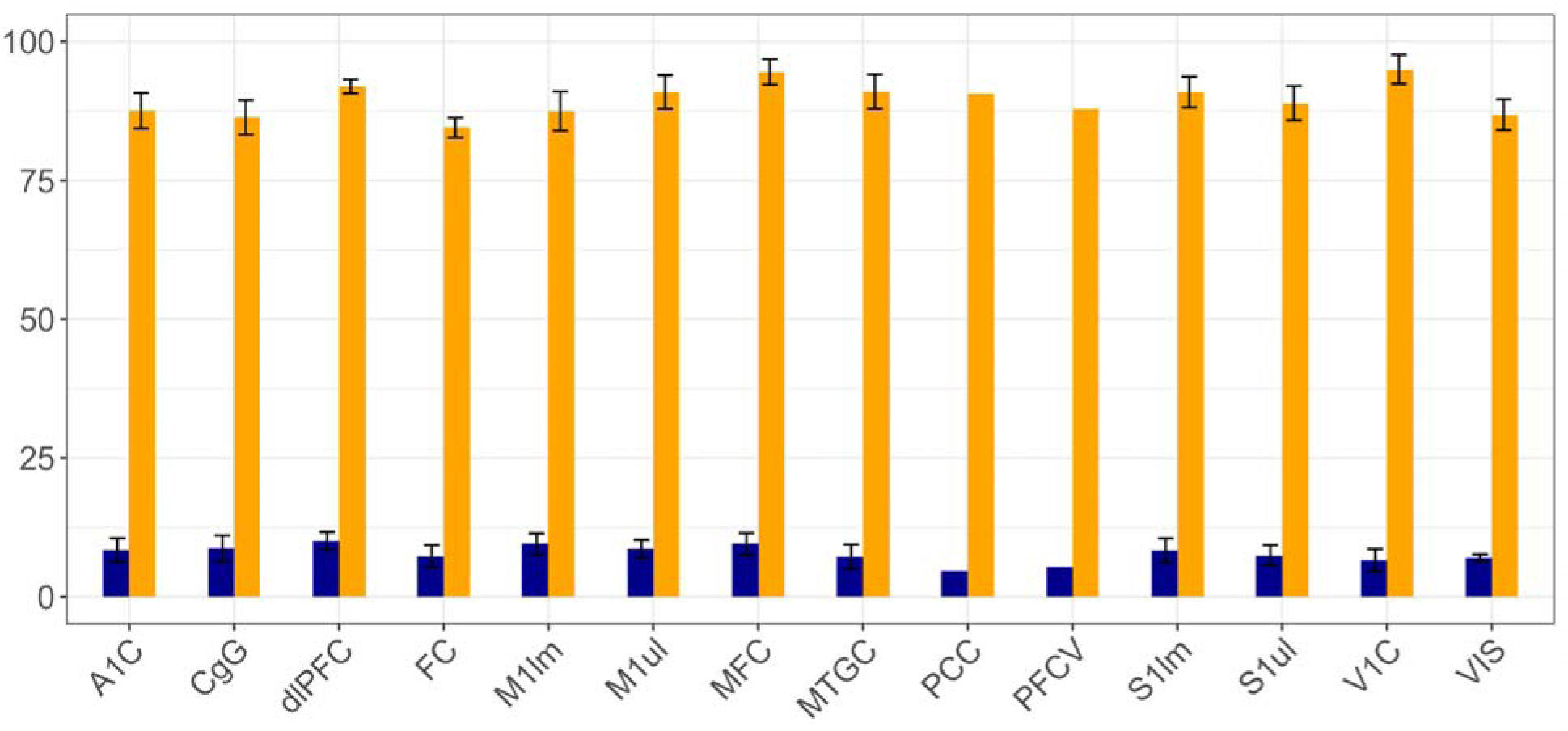
High fidelity and low background levels of OPC subtype markers calculated using our method across all studies with subtype resolution.

